# Nanopore ion sources deliver single amino acid and peptide ions directly into high vacuum

**DOI:** 10.1101/2021.08.15.456243

**Authors:** Nicholas Drachman, Mathilde LePoitevin, Hannah Szapary, Benjamin Wiener, William Maulbetsch, Derek Stein

## Abstract

We report a nanopore ion source for mass spectrometry that delivers ions directly into high vacuum from aqueous solutions. The ion source comprises a pulled quartz pipette with a sub-100 nm opening. Ions escape an electrified meniscus by ion evaporation and travel along collisionless trajectories to the ion detector. We measure mass spectra of 16 different amino acid ions, post-translationally modified variants of glutathione, and the peptide angiotensin II, showing that these analytes can be emitted as desolvated ions. The emitted current is composed of ions rather than charged droplets, and more than 90% of the current can be recovered in a distant collector. By circumventing the sample loss mechanisms inherent to conventional electrospray ionization (ESI), where charged droplets are sprayed into a background gas that scatters ions and degrades their transmission, the nanopore ion source could enable more sensitive proteomic analyses.

## 1 Introduction

Mass spectrometry (MS) is the workhorse of proteomics research thanks to its ability to distinguish amino acids and small peptides by their mass to charge ratio [1]. Its utility also significantly derives from the availability of soft ionization techniques for transferring peptide ions into the gas phase intact [2]. In particular, electropray ionization (ESI) transfers analyte into a mass spectrometer via a plume of charged droplets that emerge from a liquid cone-jet at the end of a voltage-biased emitter, as illustrated in Fig. 1a [3, 4]. The droplets pass through a background gas that induces a series of evaporation and Coulomb explosion cycles that ultimately release analyte ions into the gas phase [5]. However, the background gas needed to liberate ions from droplets is also a source of significant sample loss that limits the sensitivity of MS.

**Fig. 1.**
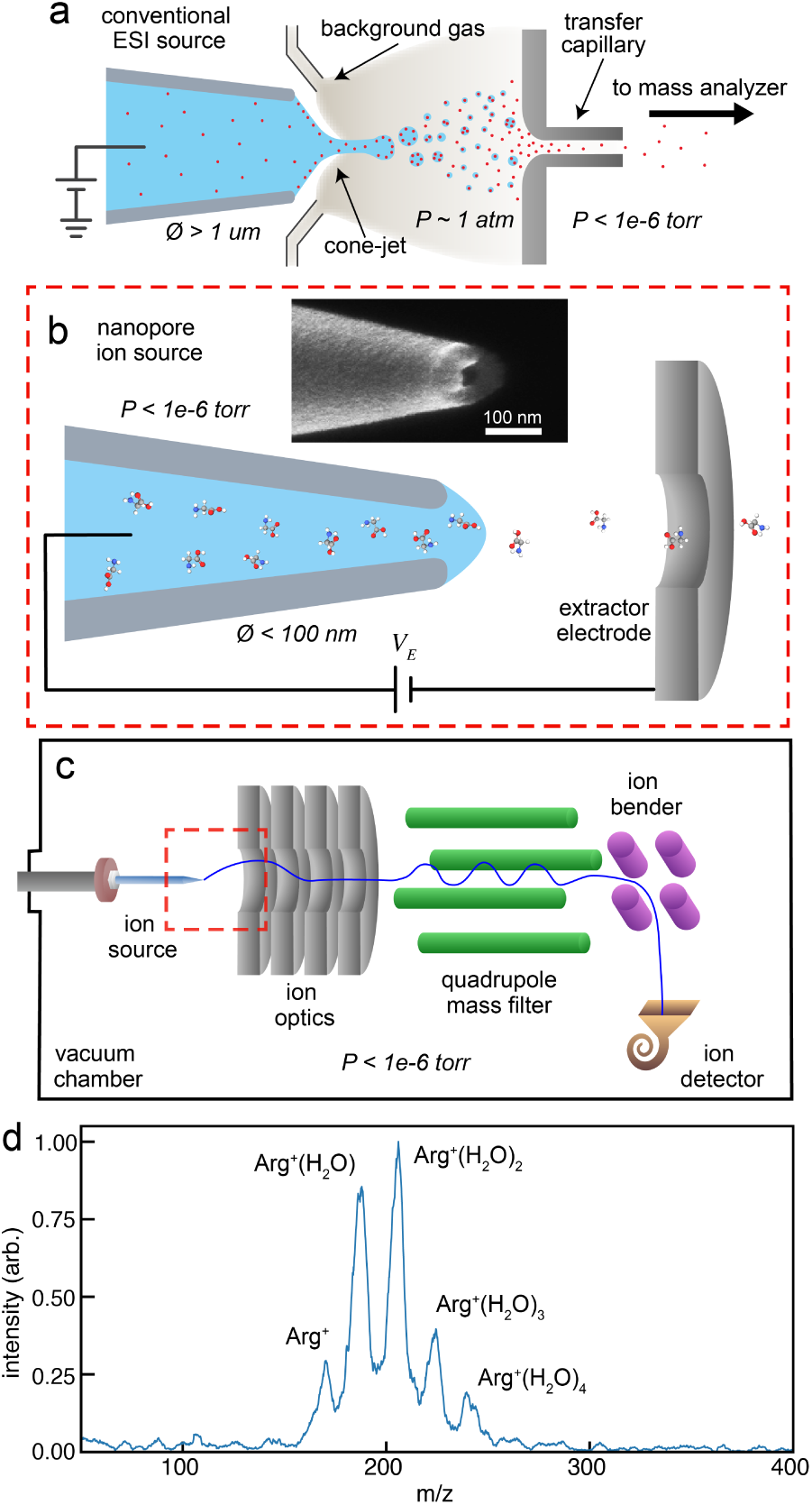
Comparison of conventional electrospray ion source with nanopore ion source. **a**, Schematic of conventional electrospray ionization highlighting the background gas that stimulates evaporation of solvent from droplets, as well as the transfer capillary where significant ion loss occurs. **b**, Schematic of a nanopore ion source showing the liquid-filled nanopipette tip, the extractor electrode, and the extraction voltage *V_E_*applied between them. Inset shows an SEM image of the tip of a pulled quartz nanopipette with a tip ID of 30 nm. **c**, Schematic of the mass spectrometer used in this study. Ion optics comprising an extractor electrode and an einzel lens extract ions from the liquid meniscus at the ion source and focus them through a quadrupole mass filter and an electrostatic ion bender. The transmitted ions strike a channel electron multiplier detector which is sensitive to single ions. **d**, Mass spectrum of 100 mM arginine in aqueous solution obtained with a 41 nm ID nanopore ion source in our quadrupole mass spectrometer.

The background gas and the plume of charged droplets it creates widely disperse ions, the majority of which collide with the transfer capillary, which bridges the ambient-pressure ion source and the first pumping stage of the mass spectrometer, and other hardware components upstream of the detector [2]. The ion transmission efficiency of an ion source, which refers to the fraction of emitted ions that enters the mass analyzer, is a key determinant of the sensitivity of mass spectrometry [6]. Early ESI sources had emitter tips with diameters of hundreds of micrometers, and only one ion in *∼*10^4^ reached the mass analyzer [7]. Nano-electrospray ionization (nano-ESI) increased the ion transmission efficiency to *∼*1% in typical measurements (occasionally reaching as high as 12%) by using emitters with micrometer-scale tips that reduced the flow rates to the range of several nL/min [8–10]. Marginean *et al.* demonstrated that by optimizing the background gas pressure, the sheath gas flow rate, and the ion funnel voltages of a subambient pressure nano-ESI source, they could achieve an ion utilization efficiency (the fraction of analyte originally in solution reaching the detector) as high as 50% for a particular small peptide [11]. However, co-optimizing the efficiency of multiple analytes is fundamentally challenging because ESI involves processes that physically separate different ion species within the plume [12]. State-of-the-art MS instrumentation still requires thousands to millions of copies of proteins for their identification [13, 14]. This sensitivity falls short of what is desired for single-cell proteomics and single-molecule analyses [15–18]. Achieving single-molecule sensitivity requires an ion source that circumvents the loss mechanisms intrinsic to spraying charged droplets into a background gas.

We have developed a nanopore ion source that emits amino acid and small peptide ions directly into high vacuum from aqueous solution at its tip (Fig. 1b). The heart of the ion source is a pulled quartz pipette with a tip whose inner diameter (ID) is smaller than 100 nm [19, 20]. We reason that the smallness of the tip will influence ion emission in three important ways: First, the surface tension of water can maintain a stable liquid-vacuum interface that supports many atmospheres of pressure when stretched across a nanoscale opening [21]. Second, the fluid flow rate, which scales as the inverse cube of the tip diameter, is too low for a stable electrospray cone-jet to form [22, 23], and this may prevent charged droplets from being emitted altogether. Third, electric fields will concentrate at a sharp, conductive tip like an electrolyte-filled nanopipette, and may reach *∼*1 V/nm at the meniscus, the approximate field needed to draw out ions at high rates by the process of ion evaporation [24, 25].

Here we characterize the emission of amino acid and small peptide ions from aqueous solutions directly into high vacuum using a nanopore ion source. We obtain mass spectra using a custom quadrupole mass spectrometer in which the nanopore ion source operates at pressures below 10*^−^*^6^ torr (Fig. 1c) [26]. Separately, we measure the transmission efficiency between the ion source and a downstream Faraday cup for both aqueous salt solutions and aqueous amino acid solutions and find it to exceed that of ESI by two orders of magnitude. We further use a magnetic sector to separate charged droplets from small ions and thereby demonstrate that the nanopore source can be made to emit only ions. We also measured the pH-dependence of amino acid ion emission rates to probe the emission mechanism.

## 2 Results

### 2.1 Emitting amino acid ions from a nanopore ion source

We characterize the emission of amino acids from aqueous solution in a custom quadrupole mass spectrometer shown in Fig. 1c [26]. In a typical experiment, ion emission from a nanopore ion source is initiated by applying an extraction voltage, *V_E_*, in the range +190 V to +380 V between the tip and the extraction electrode. The emission current, *I_E_*, is typically in the range of 2-20 pA. Significantly higher currents, ranging from 100s of pA to a few nAs are often measured when working with aqueous salt solutions. The onset of current is abrupt and accompanied by the measurement of ions striking the instrument’s detector. We collect easily interpretable mass spectra within seconds to minutes at these low emission currents. We observed that an individual nanopore ion source can be kept in operation for several weeks without clogging.

Fig. 1d shows a typical mass spectrum obtained from a 100 mM solution of arginine in water. This spectrum is obtained in positive ion mode using a nanopore ion source with inner tip diameter of 41 nm. Five peaks are clearly visible. The peak at 175 *m/z* corresponds to the singly protonated arginine ion (labelled as Arg^+^, shorthand for the protonated form [Arg + H^+^]). The higher *m/z* peaks are all separated by 18 *m/z*, the shift induced by an additional water molecule. Thus, the other peaks correspond to solvated arginine clusters (labelled as Arg^+^(H_2_O)*_n_*, where the integer *n* denotes the number of attached water molecules).

Figure 2a illustrates how the tip diameter can influence the mass spectrum obtained from an aqueous amino acid solution, again using 100 mM arginine. The spectra shown are obtained using tips with inner diameters of 20 nm, 125 nm, and 300 nm. The largest tip produces a broad spectrum of peaks that includes the bare arginine ion, eight incrementally hydrated arginine ion clusters, and a peak at 349 *m/z* that corresponds to the arginine dimer ion 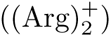. The intermediate-sized tip produces a narrower spectrum that includes the bare arginine ion, six incrementally hydrated arginine ion clusters, and a relatively diminished arginine dimer ion peak. The smallest tip primarily produces the bare arginine ion, but attenuated peaks corresponding to the singly and doubly hydrated arginine ion clusters are also visible in the spectrum. Smaller tips tend to produce relatively stronger signals and less noisy spectra than larger tips, as can be seen by comparing the baselines of the three spectra in Fig. 2a. We observe some variance in the distribution of solvation states between nanopore ion sources with similar tip sizes, and even in repeated measurements with the same tip. This is illustrated, for example, by the different distributions of arginine solvation states in Fig 2a (20 nm I.D.) and in Fig 5b (24 nm I.D.). However, only tips with inner diameters smaller than about 65 nm produced spectra where most of the amino acid ions were in the desolvated state.

**Fig. 2.**
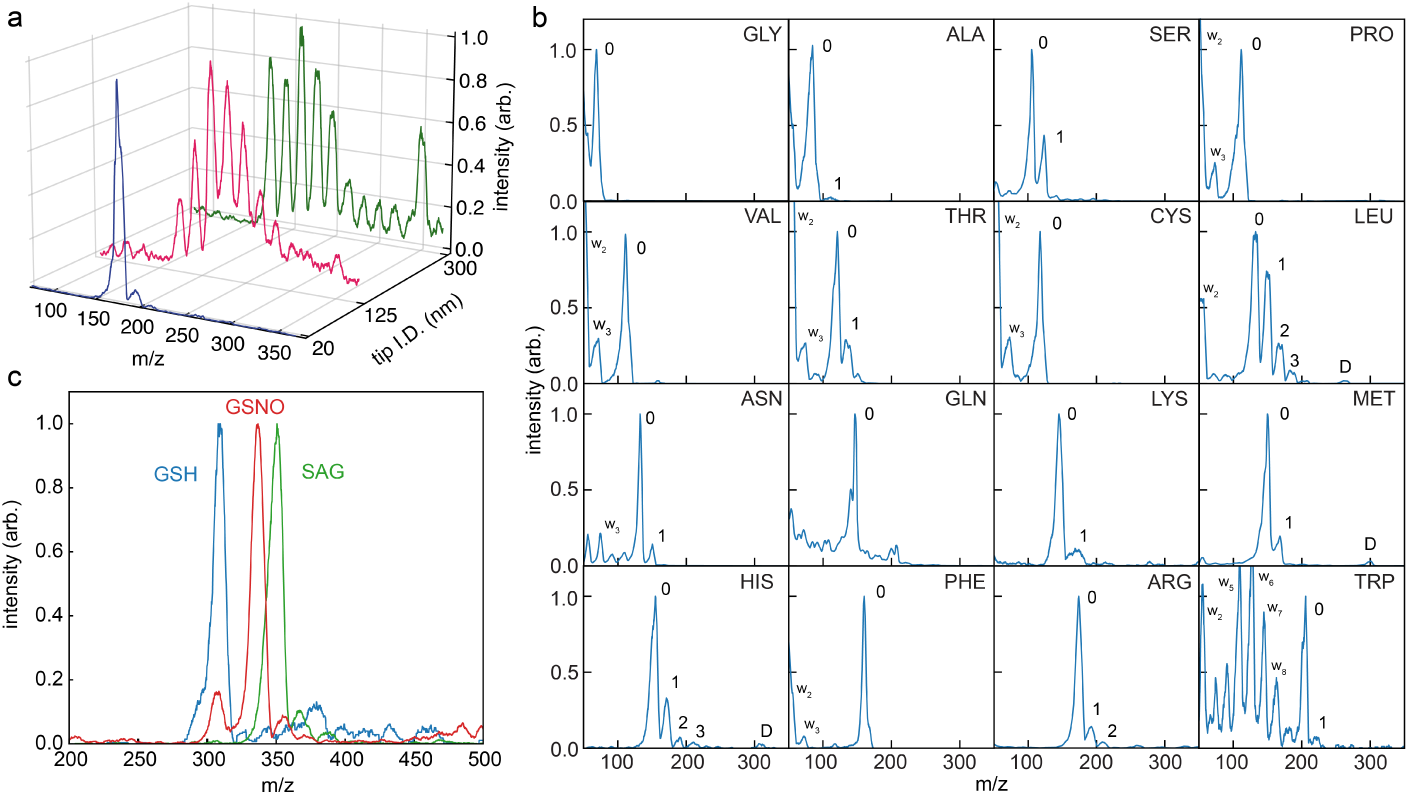
Amino acid and peptide spectra. **a**, Mass spectrum of 100mM arginine solution in H_2_O using nanopore ion sources with 3 different inner tip diameters (20 nm, 125 nm, and 300 nm). **b**, Gallery of 16 amino acid mass spectra, ordered from top left to bottom right by mass. Peaks labelled *w_n_* correspond to water clusters (H_3_O^+^(H_2_O)*_n_*), peaks labelled 0 correspond to the desolvated amino acid ion, 1 corresponds to the singly solvated amino acid ion, etc, and peaks labelled D correspond to the desolvated amino acid dimer ion. All experiments were carried out using nanopore ion sources with 20-60 nm inner tip diameters (see Supplementary Information Table 1). Each spectrum comprises thousands of individual detection events. Supplementary Fig. S2 shows how the mass spectrum develops with increasing number of events. **c**, Overlaid mass spectra of glutathione (GSH, blue) and two of its PTM variants, s-nitrosoglutathione (GSNO, red) and s-acetylglutathione (SAG, green).

Figure 2b shows mass spectra obtained from 16 different aqueous amino acid solutions, all at 100 mM concentration with the exception of tryptophan which was 50 mM. Four different tips with inner diameters of 20, 25, 57 and 58 nm were used for these measurements. The most prominent amino acid peak in every spectrum shown in Fig. 2b corresponds to a singly charged and unsolvated ion. The spectra for glycine, alanine, proline, valine, cysteine, glutamine, and phenylalanine show no additional peaks which could correspond to solvated amino acid ions. The spectra for serine, threonine, asparagine, lysine, methionine, histidine, arginine, and tryptophan show a secondary peak 18 *m/z* to the right of the unsolvated peak, corresponding to the singly hydrated amino acid ion. Leucine shows a third and fourth peak corresponding to higher solvation states. The spectra for leucine, methionine, and histidine show additional small peaks corresponding to bare, singly charged amino acid dimers. A number of spectra also show peaks below 200 *m/z* corresponding to singly protonated water clusters (H_3_O^+^(H_2_O)*_n_*); the same peaks appear in control measurements of aqueous solutions with no amino acid present (see Supplemental Information Fig. S1). Four proteinogenic amino acids are absent from Fig. 2b: we do not attempt to measure aspartic acid and glutamic acid in positive ion mode because of their low isoelectric points; we also left out isoleucine because it is indistinguishable from leucine based on *m/z*; and tyrosine gave poor emission characteristics, likely related to its low solubility in water. We worked with high (50 mM - 100 mM) analyte concentrations to obtain interpretable spectra quickly. Reliable identification of amino acids in these measurements required the release of only attomoles of analyte from the ion source, as shown in the Supplemental Information §3. The minimum concentration of analyte at which identification is possible is an alternative measure of sensitivity that we did not explore in this work.

### 2.2 Measuring post-translationally modified peptides

Figure 2c shows mass spectra of glutathione and two chemically modified variants, s-nitrosoglutathione, and s-acetylglutathione. Glutathione is a tripeptide (amino acid sequence: GSH) found in high concentrations in most cells [27], and the variants we studied result from common post-translational modifications [28, 29]. Ion sources with 20 nm ID tips generate the peptide ions from 100 mM aqueous solutions with a pH between 3.1 and 3.9, adjusted by the addition of acetic acid. The glutathione spectrum shows a single peak at 308 *m/z*, which corresponds to the singly protonated, unsolvated glutathione ion. The spectra of s-acetylglutathione and s-nitrosoglutathione show dominant peaks at 350 *m/z* and 337 *m/z*, respectively, corresponding to the singly protonated, unsolvated peptide ions; each spectrum also shows two progressively smaller peaks 18 and 36 *m/z* to the right of the dominant peak, corresponding to singly and doubly solvated peptide ions, respectively.

### 2.3 Measuring a multiply charged peptide

Figure 3 shows mass spectra obtained from an experiment where an aqueous glutathione solution was replaced with an angiotensin II solution (amino acid sequence: DRVYIHPF) inside a 29 nm ID tip. The mass spectrum of the 100 mM glutathione solution (pH 3.0) initially inside the tip shows a large peak at 308 m/z that we assign to protonated glutathione, along with additional smaller peaks at 326, 344, and 615 m/z that we assign, respectively, to singly protonated, singly hydrated glutathione, singly protonated, doubly hydrated glutathione, and singly protonated glutathione dimer. We gradually introduced a 1 mM angiotensin II solution into the tip using a previously described tube-in-tube fluid handling system [26]. Additional peaks appeared in the mass spectrum at 524 m/z and 350 m/z, which we assign to doubly and triply protonated angiotensin II, respectively. The emission current throughout the measurement was 10 *±* 3 pA with an extraction voltage of 400 V.

**Fig. 3.**
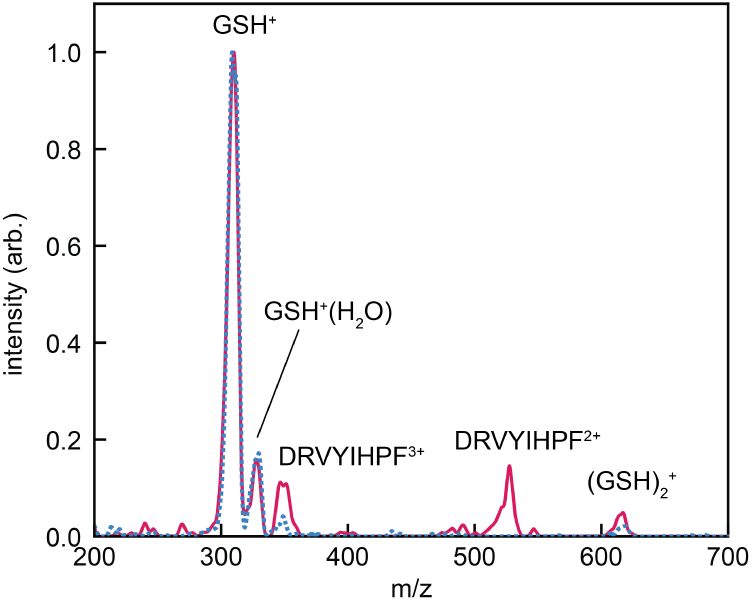
Mass spectra of 100 mM glutathione (dashed blue line) and a mixture of glutathione and angiotensin II obtained about 45 minutes after introducing a 1 mM angiotensin II solution into the tip of the ion source (solid red line). Peaks assigned to glutathione (GSH^+^), singly hydrated glutathione (GSH^+^)+H_2_O)), glutathione dimer 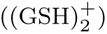, and the doubly and triply protonated states angiotensin II (DRVYIHPF*^n^*^+^, *n* = 2, 3) are indicated.

### 2.4 Ion transmission efficiency

We measure the efficiency with which ions pass from the nanopore source to a distant detector in a high vacuum environment as illustrated in Fig. 4a. Ions emitted by the source are focused into the 2.4 cm diameter opening of a Faraday cup located *∼* 50 cm away. The ion transmission efficiency is the ratio of the current transmitted to the Faraday cup, *I_T_* , to the current emitted from the nanopore, *I_E_*. Figure 4b shows *I_E_*, *I_T_* , and the ion transmission efficiency measured over several minutes using a 39 nm ID tip filled with a 100 mM aqueous solution of sodium iodide. The mean ion transmission efficiency in this measurement is 93.4 % *±* 1.7 %. Figure 4c shows the results of a similar measurement made with a 49 nm ID tip filled with 100 mM arginine in aqueous solution at pH 8.95. The mean ion transmission efficiency is 65.3 % *±* 8.1 %. In both measurements, small changes in *I_E_* are mirrored by *I_T_* . We had more difficulty optimizing the focusing voltages in the amino acid measurement because of the very low emission currents, which were at the single pA level.

**Fig. 4.**
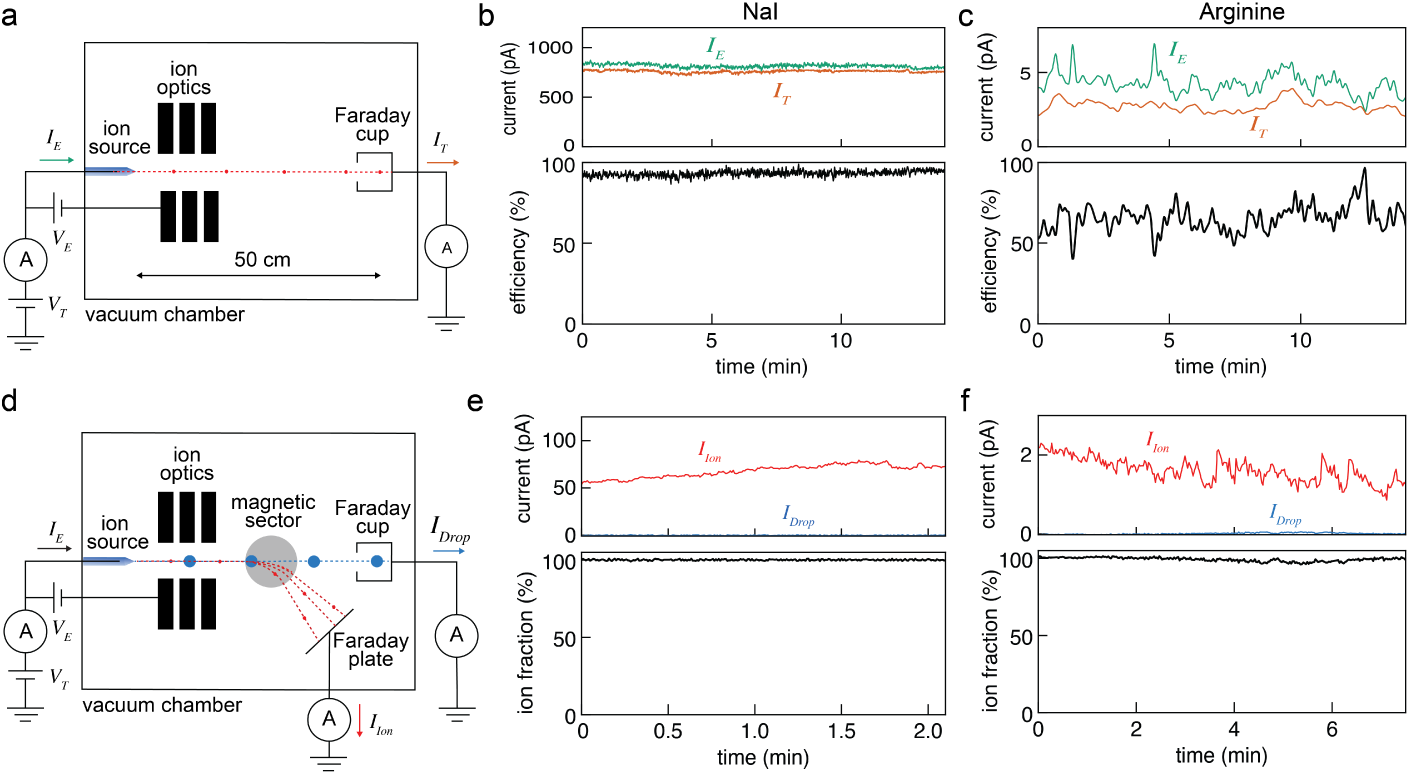
Ion transmission efficiency and ion/droplet current fractions. **a**, Setup used to measure the ion transmission efficiency of a nanopore ion source. **b**, *I_E_* and *I_T_* (above) and the resulting transmission efficiency (below) of a 39 nm ID nanopore ion source containing a 100 mM aqueous solution of NaI. **c**, *I_E_* and *I_T_*(above) and the resulting transmission efficiency (below) of a 49 nm ID nanopore ion source containing a 100 mM aqueous solution of arginine at pH 8.95. **d**, Setup used to separately measure *I_Ion_*and *I_Drop_*. **e**, *I*_Ion_ and *I*_Drop_ (above) and the resulting ion fraction (below) emitted by a 39 nm ID nanopore ion source containing a 100 mM aqueous solution of NaI. **f**, *I*_Ion_ and *I*_Drop_ (above) and the resulting ion current fraction (below) emitted by a 41 nm ID nanopore ion source containing a 100 mM aqueous solution of Arginine at pH 8.95.

**Fig. 5.**
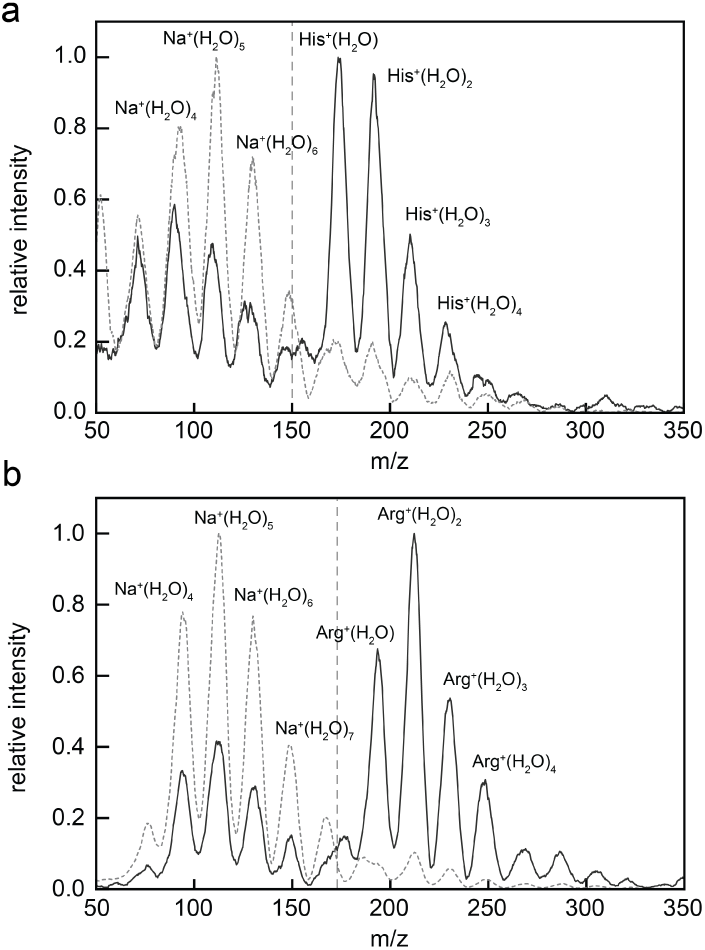
Amino acid ion emission pH. **a** Normalized mass spectra of 100 mM Histidine and 24 mM NaI/NaOH in aqueous solution at pH 5.49 (darker solid curve) and 8.52 (lighter dashed curve) respectively. **b** Normalized mass spectra of 100 mM Arginine and 50mM NaI at pH 9.10 (darker curve) and 10.78 (lighter curve) respectively. The vertical dotted line shows the m/z value used in the analysis to separate arginine signal from sodium signal. In both panels, the vertical dashed line shows the m/z value used in the analysis to separate the amino acid signal from sodium signal.

### 2.5 Separating ions and charged droplets

We investigate the possibility that the nanopore ion source emits charged droplets in addition to ions by adding a magnetic sector to the flight path as shown in Fig. 4d. The 6 cm diameter, 0.54 T magnetic sector deflects charged species based on their mass-to-charge ratio. Droplets with an *m/z >* 33, 000 deflect by less than 3.1*^◦^* and enter the Faraday cup where their current, *I*_Drop_, is measured. We do not expect the droplet size to change appreciably between emission and detection because nanoscale aqueous droplets freeze rapidly in vacuum [30, 31]. Meanwhile, ions with m/z in the range of about 70 to 325 deflect onto a separate Faraday plate where the ion current, *I*_Ion_, is measured (see Supplementary Information §5). Figure 4e shows *I*_Ion_, *I*_Drop_, and the ion fraction, defined as 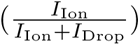, for a measurement performed using a 28 nm ID tip filled with a 100 mM aqueous solution of NaI. *I*_Ion_ varies between about 60 pA and 80 pA while no *I*_Drop_ is observed. Figure 4f shows the results of a similar measurement made using a 41 nm ID tip filled with a 100 mM aqueous solution of arginine at pH 8.75. *I*_Ion_ varies between about 1 pA and 2 pA while there is no detectable *I*_Drop_. In both measurements, the nanopore source was only observed to emit ions.

### 2.6 pH dependence of amino acid emission

To further investigate the emission mechanism, we measured the pH-dependence of the emission rates of histidine and arginine. For each amino acid, we prepared solutions at two different pH values but with a fixed concentration of Na^+^ to serve as a reference ion. We made all measurements with a single, 24 nm ID tip. Figure 5a shows the mass spectra of 100 mM aqueous histidine solutions with pH values of 5.49 and pH 8.52; the concentration of Na^+^ is 24 mM. At pH 5.49, peaks corresponding to the bare histidine ion and several solvated states appear between approximately 150 m/z and 275 m/z; the histidine peaks are more intense than the solvated sodium peaks which appear between approximately 50 m/z and 150 m/z. At pH 8.97, the histidine peaks are still visible, but their intensity is significantly diminished relative to the sodium peaks. Figure 5b shows the mass spectra of 100 mM arginine solutions with pH values of 9.10 and 10.78; the solutions also contain 50 mM of Na^+^. Similarly to histidine, bare and solvated arginine peaks, which appear in the range between 170 m/z and 350 /mz, have a higher intensity than the sodium peaks at the lower pH of 9.10. The relative intensity of the arginine peaks decreases dramatically at the higher pH of 10.78. The results shown in Fig. 5 are representative of repeated measurements. To quantify the pH-dependence of the amino acid emission rates, we normalized them by the sodium emission rates. The histidine emission rate increased by an average factor of 8.14 *±* 1.65 as the pH decreased from 8.52 to 5.49. The arginine emission rate increased by an average factor of 22.6 *±* 10.5 as the pH decreased from 10.78 to 9.10.

### 2.7 Calculating the probability of ions scattering

Calculations indicate that most ions trace collisionless trajectories from the ion source to the detector. Figure 6a shows the probability that an amino acid ion dressed with a hydration shell will collide with a gas molecule, based on the kinetic theory of gases. We assume that ions pass through a distribution of evaporating water molecules [32] and a homogeneous low-pressure background of N_2_. Figure 6a sketches the physical situation and plots the number density of gas molecules and the cumulative collision probability as functions of the distance from the meniscus. The cumulative probability of an ion colliding with a gas molecule over the entire 50 cm trajectory from the source to the detector is just 2.1%, meaning the vast majority of ions do not experience any collisions. Most of the collisions occur within 200 nm of the liquid meniscus due to the high density of evaporated water molecules there. A detailed description of these calculations is found in §4 of the Supplementary Information.

**Fig. 6.**
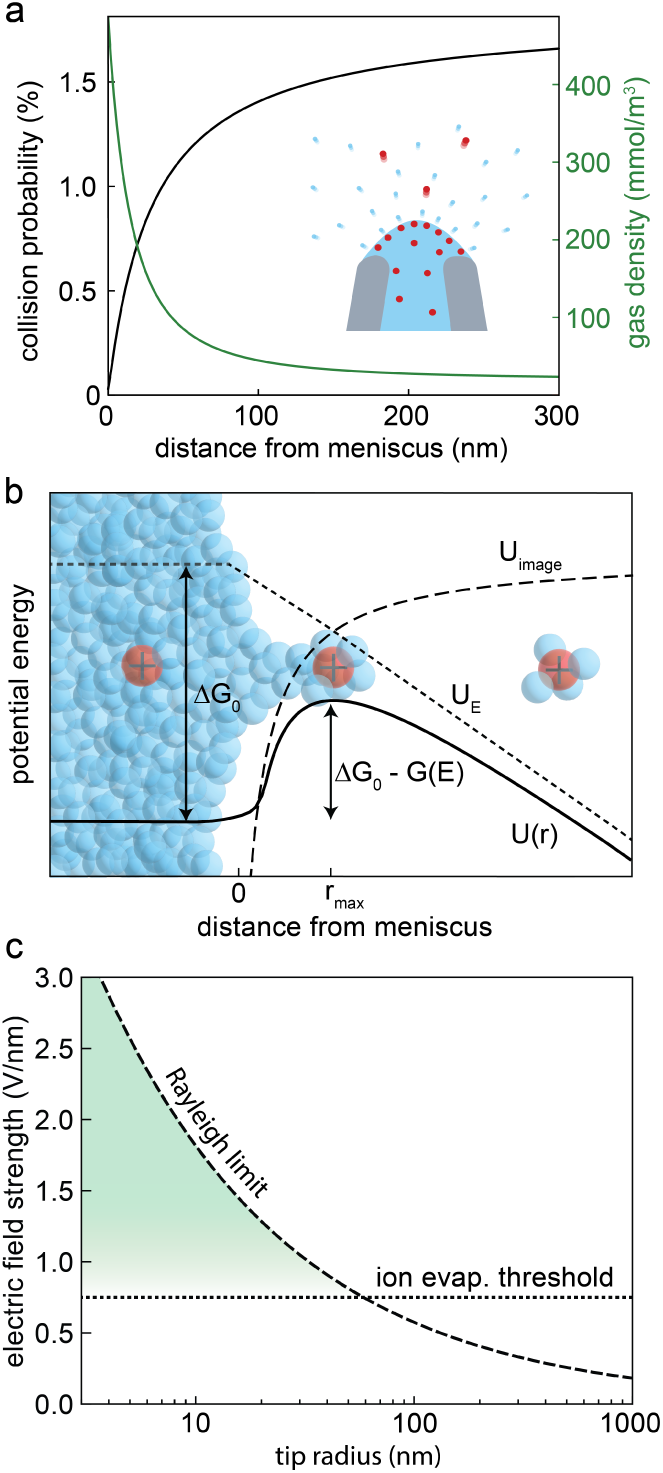
Ion emission mechanism. **a** Plot of the cumulative probability that an emitted amino acid with a small hydration shell (radius = 7 Å) will collide with an evaporated water molecule or background gas molecule as a function of distance from the meniscus, *r*. The green curve shows the calculated maximum water vapor density as a function of the distance from the meniscus. **b** Potential energy diagram showing the contributions to the energy barrier for ion evaporation: the solvation energy, Δ*G*_0_, image charge attraction, *U*_image_, and electrostatic repulsion, *U*_E_. **c** Plot of the approximate limiting fields for ion evaporation and droplet generation (Rayleigh limit). The region where ion evaporation should dominate is highlighted in green.

## 3 Discussion

There are two distinct mechanisms by which the nanopore source could generate the ions that we measure. The first is conventional electrospray ionization, a multi-step mechanism in which an ion source first produces charged droplets which go on to evaporate ions into the vapor phase [2]. The second is a single-step mechanism in which ions evaporate directly from the electrified meniscus of the ion source [33]. We highlight the important physical distinctions between these two mechanisms before presenting an analysis that rules out the former.

ESI begins when a voltage applied to the liquid sample generates electrical forces on the meniscus that are strong enough to overcome the surface tension and draw out charged droplets (Fig. 1a). The droplets typically measure hundreds of nanometers to tens of micrometers in radius and contain fully solvated analyte molecules [34]. A background gas is used to evaporate solvent from the droplets [34]. As a droplet shrinks, its charge density increases, and the resulting internal Coulomb forces can once again overcome surface tension when the electric field strength at the surface reaches the Rayleigh limit, 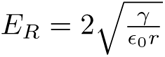, where *γ* is the surface tension, *ɛ*_0_ is the permittivity of free space, and *r* is the droplet radius [3, 35]. The unstable droplet undergoes a Coulomb explosion and expels smaller charged droplets [3, 5]. The cycle repeats, producing ever smaller droplets requiring ever stronger electric fields to reach the Rayleigh limit. When the field strength at the surface reaches about 10^9^ V/m, it becomes possible for ions to quickly escape the droplet by a different process: Ion evaporation is a thermal process in which ions overcome an energy barrier associated with their solvation energy, 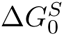, to enter the vapor phase [24]. Coulomb explosions cease and ion evaporation becomes the main charge release mechanism for small molecules when droplets reach around 10 nm in radius [24, 36, 37].

Alternatively, the nanopore source might deliver ions and ion clusters into the vapor phase directly from the meniscus at the tip by an ion evaporation process. This alternative mechanism circumvents the emission of droplets and their interactions with a background gas that are integral to ESI. Figure 6b illustrates the direct evaporation of a solvated ion cluster from an electrified meniscus and a schematic of the energy landscape [24, 38]. An ion inside the nanopore source is fully solvated by water molecules, which reduce the free energy by an amount 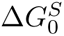, called the solvation energy. The free energy increases as the ion protrudes from the meniscus, distorting it against surface tension. At this point, the ion is also attracted by its own image charge as it begins to break free of the solvent. Far from the meniscus, Coulomb forces drive the ion (or cluster) away from the tip and its free energy decreases. Modelling the height of the energy barrier as 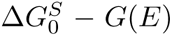, where *G*(*E*) is the field-dependent reduction in the barrier height [3, 24, 38], the current density of ions escaping the meniscus is given by

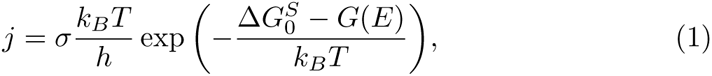

where *σ* is the surface charge density, *k_B_T* is the thermal energy, and *h* is the Planck constant. The ion evaporation current density only becomes appreciable when the electric field at the meniscus approaches *∼* 10^9^ V/m, at which point 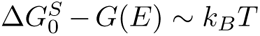 [39]. The ion evaporation mechanism applies equally well to bare ions and small ion clusters, as each is a charged species that can escape the liquid over a solvation energy barrier [24, 40, 41]. On the other hand, charged nano-droplets comprising hundreds of solvent molecules or more are too large to evaporate.

Multiple lines of evidence lead us to conclude that the nanopore source produces ions directly by ion evaporation rather than by the conventional ESI mechanism in which ions emerge from charged droplets. First, the chain of Coulomb explosions that characterizes ESI cannot occur under the high vacuum conditions of our experiments. Droplets quickly freeze before shedding a significant fraction of their mass when there is no background gas to provide the thermal energy that normally sustains solvent evaporation [30]. A previous study showed that nanoscale charged aqueous droplets survive under high vacuum for more than a second without undergoing Coulomb explosions [42].

Secondly, we searched for droplets and measured none among the charged species delivered by the source (Fig. 4d-f). Our measurements, which are sensitive to species with *m/z >* 33, 000, exclude droplets with a radius larger than 14 nm and a charge as high as the Rayleigh limit. For comparison, nano-ESI begins with a spray of droplets that are in the detectable range of tens to hundreds of nanometers in diameter [2, 43]. If even smaller charged droplets were produced [44, 45], they would evaporate ions readily, lose a significant fraction of their charge, and move into the detectable range of *m/z*; when the electric field at the surface falls to 0.75 V/m, the approximate threshold for ion evaporation, droplets as small as 7 nm in radius are detectable.

Thirdly, the nanopore source is sufficiently small that as the electric field at the liquid meniscus increases, it reaches the threshold for ion evaporation well before the Rayleigh limit. Figure 6c, which compares the critical electric field for ion evaporation with the Rayleigh limit, illustrates why ions evaporate from the meniscus while the meniscus remains stable. If we consider a nanopore source with a radius of 15 nm, we expect ions to evaporate from the meniscus when the electric field reaches about 0.75 V/nm, whereas droplets cannot emerge unless the electric field grows about twice as strong. On the other hand, a conventional ESI source with a radius of 1 *µ*m produces charged droplets when the electric field at the meniscus reaches about 0.2 V/nm, which is well below the threshold for ion evaporation. The crossover from a conventional droplet-emitting source to a nanopore source that can evaporate ions directly is predicted to occur at a radius of about 60 nm.

Finally, the pH-dependence of the emission rates of arginine and histidine we measured (fig. 5) is consistent with a simple kinetic model of ion evaporation from the meniscus. We consider a two-step process in which an analyte molecule M can undergo reversible protonation/deprotonation with rates *k*_on_[H^+^] & *k*_off_, and non-reversible evaporation from the protonated state with rate *k*_evap_,

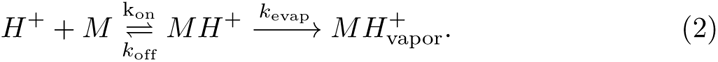

We assume that evaporation is the rate-limiting step and that *k*_evap_ *≪ k*_off_, justified by the picoampere-level currents measured in this work. Holding the concentration [MH^+^] constant, we find that the quasi-steady-state emission rate, *v ≡* [*MH*^+^]*k*_evap_, scales as

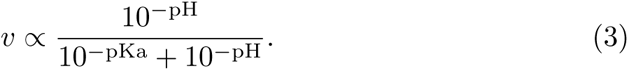

Figure 7 plots the relative emission rates predicted by this model for both histidine and arginine and compares them to the measured values. For histidine, two protonation reactions occur within the experimental pH range (with pKa_1_ = 6.04 and pKa_2_ = 9.09); a single dissociation constant corresponding to the isoelectric point (pKa=7.59) models the combined effect [46]. For arginine, only the protonation of the amino group (pKa= 9.00) is relevant [46, 47]. The kinetic model predicts that the emission rates should increase by factors of 9.2 and 27.3 over the pH ranges investigated for histidine and arginine, respectively, in agreement with our measured values of 8.14 *±* 1.65 and 22.5 *±* 10.5. By contrast, ESI emits arginine and histidine at rates that depend much more weakly on pH (by a factor less than three over the same pH range), and this observation was attributed to the acidification of droplets that accompanies the ESI process [48, 49].

**Fig. 7.**
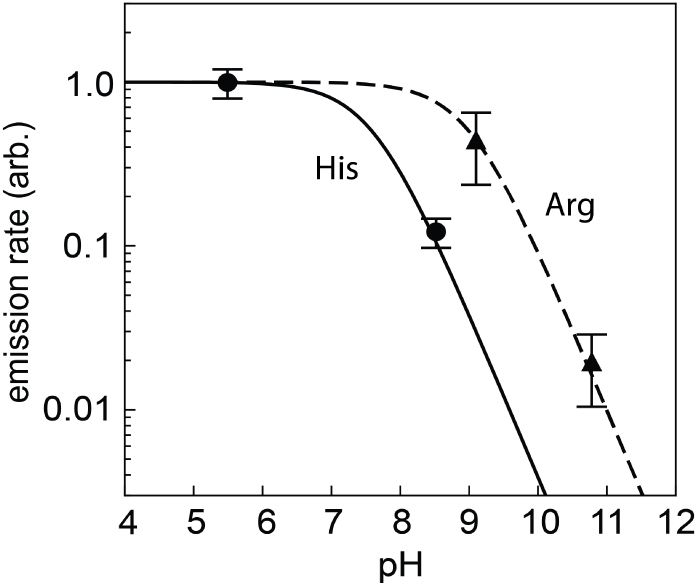
Amino acid ionization versus pH. Summary of repeated measurements of the relative emission rates of histidine (circles) and arginine (triangles) compared to theoretical predictions. The emission rate at the lower pH, *v*_low_, is calculated using Eq. (3), and the emission rate at the higher pH is calculated using *v*_low_*/F* , where *F* is the ratio of measured signal between low and high pH. The error bars represent standard error of the mean.

The nanopore source appears to operate in a pure ion mode where no droplets are emitted. Pure ion emission was previously observed from highly conductive liquids including liquid metals [50], ionic liquids [33, 51], and concentrated electrolyte solutions in formamide [39, 40], but not aqueous solutions. Previous attempts to evaporate ions directly into vacuum from a fixed water meniscus encountered significant challenges, as the evaporation of water led to electrical arcing and freezing [52]. The nanopore source overcomes those challenges because the evaporation rate from the nanoscale meniscus is minuscule (less than 1 pL of water evaporating per minute); this low gas load also eases the pumping requirements of the instrument [26]. Previously, even the smallest ion emitters were used in combination with a background gas and the emission mechanism was presumed to be conventional ESI [53].

It is striking that we measure mostly desolvated amino acid and small peptide ions in many experiments (fig. 2b and c, fig. 3). This property simplifies the interpretation of mass spectra without requiring a drying gas. Were the ions emitted in an already desolvated state, or clustered with one or more water molecules which were shed on the way to the detector? Given that just 2% of emitted ions collide with a gas molecule even once (see Fig. 6a), post-emission collisions can be dismissed as a primary desolvation mechanism. Furthermore, the tip size appears to influence the solvation state, as only sub- 100 nm nanopores produced bare ions (Fig. 2a). These observations suggest that it is the local environment at the source that controls the solvation state rather than processes occurring in flight. The conditions that give rise to bare ion emission are not entirely clear. The electric fields that develop near the smallest tips are strong enough to reorient and weaken hydration bonds, which may play a role [54, 55]. Joule heating may also play a role by raising the local temperature and further weakening hydration bonds [56, 57]. In addition to single amino acid ions, we observed desolvated amino acid dimer ions in some spectra (e.g., Fig. 2b, His). This is likely due to the aggregation of amino acids in solution, which is known to occur at concentrations of 1 mM and above [58]. We also observed glutathione and its post-translationally modified variants as bare, singly charged ions. Subsequent work should investigate how the charge and hydration states depend on the peptide length, composition, and other properties.

The present study makes three significant contributions: First, our measurements and analyses of the nanopore source provide strong evidence for the direct ion evaporation mechanism. Second, the nanopore source achieves near unit ion transmission efficiency. This holds the promise of significant improvements in the sensitivity of MS measurements, which are universally desirable and could be achieved by combining a nanopore source with a sensitive mass filter (e.g., a trap-based mass filter). Third, the nanopore source is compatible with biologically relevant aqueous samples.

The findings reported here open new avenues for research. We envision sequencing a single protein molecule by fragmenting it and delivering its constituent amino acids into a mass spectrometer in sequential order [18, 26, 59]. The nanopore ion source provides the crucial means of transmitting virtually every amino acid to the detector, and because they avoid collisions in flight, the timing of detection events can be used to accurately reconstruct the order in which they left the source. However, significant challenges persist. A means of reliably fragmenting polypeptides into free monomers is required, as is a means of preserving the order of monomers prior to emission, perhaps using force gradients [59] or a narrow nanotube emitter [60] to prevent amino acids from diffusing past one another. These findings could also impact various fields outside of single-molecule sequencing or proteomics. Electrospray propulsion, a method for generating thrust through ion emission, is currently constrained by a limited selection of non-volatile ionic liquids that can be used as propellants [61, 62], but the nanopore ion source highlights the possibility of using volatile liquids. Atom probe tomography, a technique for generating compositional maps of materials using field-emitted ions, is entering a new frontier with studies of liquid samples [63–65], which may be simplified by using a nanopore ion source. Finally, the nanopore source may shed new experimental light on the ionization of water, its orientation, or its evaporation at highly electrified surfaces [66, 67].

## Methods

### Preparing nanopore ion sources

We pulled nanopipettes from 7.5 cm long quartz capillaries with 0.7 mm inner diameter and 1 mm outer diameter (QF100-70-7.5, Sutter Instruments) using a laser puller (P-2000, Sutter Instruments) according to the following single-line recipe: heat = 650, velocity = 45, delay = 175, pull = 190. To prepare a nanopore for imaging, we deposited approximately 5 nm of carbon to the nanopipettes using thermal evaporation (CED 030, BalTec) with double-stranded carbon thread (Leica Microsystems). We imaged the nanopipettes by scanning electron microscopy (LEO 1530 VP, Zeiss inc.) to measure the inner and outer tip diameters. Nanopipettes were plasma cleaned in air for two minutes (Plasma Preen II-973, Plasmatic Systems Inc.) to make the surfaces hydrophilic prior to filling with sample solution.

### Amino acid solutions

Amino acid solutions comprised the amino acid of interest (Sigma-aldrich) dissolved in deionized water (18 MΩcm, Millipore) at a concentration of 100 mM, with the exception of tryptophan, whose concentration concentration was 50 mM. We reduced the pH below the amino acid’s isoelectric point by adding 0.1-0.5% v/v glacial acetic acid (Sigma-Aldrich). Peptide solutions comprised the peptide of interest (glutathione, s-acetylglutathione, or s-nitrosoglutathione) dissolved in deionized water at a concentration of 100 mM. We purchased glutathione and s-acetylglutathione in powder form (Sigma-aldrich), and synthesized S-nitrosoglutathione from glutathione according to a protocol from T.W. Hart [68]. We measured the pH and conductivities of each solution using a pH meter (Ultrabasic Benchtop, Denver Instruments) and conductivity meter (Sension+ EC71 GLP, Hach), respectively.

### The nanopore mass spectrometer

A custom-built quadrupole mass spectrometer, described previously in Bush et al [26], measured all the amino acid and peptide mass spectra presented here. The instrument, illustrated schematically in Fig. 1c, comprises a custom einzel lens, a quadrupole mass filter (QC MAX-500, Extrel inc.), an ion bender (Extrel inc.), and a channel electron multiplier detector with a conversion dynode (DeTech 413) sensitive to single ions. The quadrupole is driven by a 440 kHz RF oscillator and scans across a mass range of 50 m/z to 1000 m/z in about 0.6 s. The quadrupole’s Δm and Δres parameters were periodically tuned to optimize the peak shape while acquiring spectra of NaI in formamide. The pole bias was set to ensure the ions had a kinetic energy of about 5-10 eV as they traversed the mass filter. The base pressure of the instrument is 10*^−^*^8^ torr. A tube-in-tube system (previously described in Bush *et al* [26]) enables us to deliver fresh sample solution to the nanopipette tip while flushing away old solution without breaking the vacuum. A 150 *µ*m i.d., 360 *µ*m o.d. PEEK tube (IDEX Health and Science) carries fresh sample solution into the tip, displacing and pushing out old solution through a wider outer PEEK tube (0.04” i.d., 1/16” o.d., IDEX Health and Science). We prepare Ag/AgCl electrodes by placing a length of 0.5 mm diameter silver wire (Alfa-Aesar) in bleach for 15-20 minutes. The Ag/AgCl electrode is threaded far into the outer PEEK tubing, enabling us to control the voltage of the solution near the tip of the ion source.

### Measurements of amino acid and glutathione mass spectra

We prefilled the nanopipette with the amino acid or peptide solution using a Microfil flexible needle (World Precision Instruments) before mounting the nanopipette inserting them into the mass spectrometer. We continuously refreshed the solution at the tip by pumping solution through the tube-in-tube system at rate of 0.4 mL/hour using a syringe pump (NE-300, New Era Pump Systems). Note that this is not the flow rate of fluid emitted from the tip, as the tube-in-tube system carries the vast majority of the sample solution out of the tip through the rear, as described in Bush *et al* [26]. A voltage of +100 V or +200 V was applied to the electrode in contact with the solution inside the nanopore ion source using a high voltage source measure unit (2657A, Keithley Instruments) while a negative voltage applied to the extraction electrode was slowly strengthened using a high voltage power supply (Bertan 230, Spellman HV) until ionization was observed. The onset of emission typically occurred when *V_E_* was between 200 and 350 V.

### Ion transmission efficiency measurements

We measured ion transmission efficiencies in a custom vacuum chamber containing a nanopore ion source and a Faraday cup (Fig. 4a). A high voltage source measure unit (Keithley 2657a or Keithley 2410) measured the current emitted from the nanopore, *I_E_*, while also applying a voltage, *V_T_* , to it. A current-preamplifier (SR570, Stanford Research Systems) converted *I_T_* at the Faraday cup into a voltage that was recorded by a DAQ (NI PCIe-6251, National instruments). An 8-channel high-voltage power supply (DT8033, CAEN) supplied the focusing voltages to the ion optics. A custom Labview (National Instruments) program controlled *V_T_* and recorded *I_E_* and *I_T_* . The currents emitted from aqueous salt solutions (Fig. 4b), which ranged from 100 pA to 1 nA, were measured with a Keithley 2410. To reliably measure the low, picoampere-level currents emitted from aqueous amino acid solutions (Fig. 4c), we used a more sensitive source measure unit (Keithley 2657a) and a shielded, tri-axial cable that more effectively suppressed leakage currents. To further suppress noise, the source measure unit averaged three sequential measurements of *I_E_* with an NPLC setting of 5, then an 8*^th^*-order, low-pass Bessel filter with a cutoff frequency of 0.1 Hz filtered the data to obtain the results plotted in Fig. 4c. *I_T_*was sampled at a rate of 50 kHz, and 10k samples were averaged to give each data point in Fig. 4c. The setup records *I_E_* and *I_T_* data points sequentially with an overall sampling rate of *∼* 0.6 Hz. A small leakage current measured at the source measure unit with *V_E_* = 0 V and *V_T_* = 500 V was subtracted from measurements of *I_E_*. The leakage current was about 5 pA when using the Keihtley 2657a (arginine measurements) and about 100 pA when using the Keithley 2410 (NaI measurements) for the NaI efficiency measurement. An instrumental offset of about 200 fA was subtracted from measurements of *I_T_* .

### Magnetic sector measurements

We constructed a simple magnetic sector mass spectrometer by adding a magnetic sector and a Faraday plate downstream of the ion source in the vacuum chamber described above, as illustrated in Fig. 4d. The magnetic sector comprises a neodymium magnet and a yoke made of low-carbon magnetic iron (ASTM A848) which concentrate magnetic fields of 0.54 *±* 0.02 T in a flat circular region 6 cm in diameter and 1 cm in height. The Faraday plate is a stainless steel disk, 5 inches in diameter, mounted about 25 cm from the center of the magnetic sector, 45*^◦^* from the undeflected axis of the ions. Two separate current pre-amplifiers (Stanford Research Systems SR570) measured the currents striking the Faraday plate and cup. The currents were measured and processed in the same manner described above for the ion transmission efficiency measurements.

## Data availability statement

The data and analysis code used in this study are available at the Harvard Dataverse as Drachman, Nicholas; LePoitevin, Mathilde; Szapary, Hannah; Wiener, Benjamin; Maulbetsch, William; Stein, Derek, 2024, ”Replication Data for: Nanopore ion sources deliver single amino acid and peptide ions directly into high vacuum”, https://doi.org/10.7910/DVN/XQG6NJ, Harvard Dataverse.

## Acknowledgements

We thank Oxford Nanopore Technologies LTD. for supporting this work. Aspects of this work were also supported by the Institute for Molecular and Nanoscale Innovation at Brown University. N.D. was partially supported by the Brown IMSD program (NIH R25GM083270) and Graduate Fellowships for STEM Diversity (GFSD). H.S. was partially supported by the Brown UTRA program. B.W. was partially supported by NSF 1409577.

## Author Contributions

Conceptualization & supervision: D.S.; Research design: D.S., M.L., W.M.; Mass spectrometry measurements: M.L., N.D., and H.S.; Ion source characterization: N.D., W.M., and B.W.; Manuscript writing and editing: N.D. and D.S.; Figure preparation: N.D.; Funding acquisition: D.S.; All authors contributed to the discussions.

