## Supplemental Information for "Nanopore ion sources deliver single amino acid and peptide ions directly into high vacuum"

June 5, 2024

#### 1 Single amino acid ion measurement conditions

Table S1 displays the relevant experimental conditions of each amino acid measurement presented in Figure 2b of the main text. The table lists the solution properties (amino acid identity and mass, concentration, pH, and conductivity), the ion source properties (tip inner diameter, tip outer diameter, and tip number), as well as the time-averaged chamber pressure, extraction voltage, and emission current.

**Table S1:** Experimental conditions of amino acid measurements presented in Fig. 2b of the main text.

| Amino Acid | Mass | Conc. (mM) | pH | K (S/m) | Tip ID (nm)[a] | Tip OD (nm)[a] | P (torr)[b] | $V_e$ (V) | $I_e$ (pA) | Tip number[c] |
|---|---|---|---|---|---|---|---|---|---|---|
| Arginine | 174 | 100 | 8.06 | 0.205 | 20 | 64 | $8 \times 10^{-8}$ | 225 | 2 | 1 |
| Lysine | 146 | 100 | 5.75 | 0.486 | 20 | 64 | $5 \times 10^{-8}$ | 225 | 1 | 1 |
| Histidine | 155 | 100 | 6.22 | 0.247 | 20 | 64 | $1.3 \times 10^{-7}$ | 280 | 2 | 1 |
| Glycine | 75 | 100 | 4.96 | $1.96 \times 10^{-2}$ | 25 | 58 | $6 \times 10^{-8}$ | 190 | 4 | 2 |
| Alanine | 89 | 100 | 4.03 | $1.94 \times 10^{-2}$ | 25 | 58 | $6 \times 10^{-8}$ | 190 | 4 | 2 |
| Proline | 115 | 100 | 3.96 | $1.25 \times 10^{-2}$ | 25 | 58 | $6 \times 10^{-8}$ | 190 | 4 | 2 |
| Valine | 117 | 100 | 3.95 | $1.75 \times 10^{-2}$ | 25 | 58 | $6 \times 10^{-8}$ | 190 | 4 | 2 |
| Threonine | 119 | 100 | 3.88 | $1.82 \times 10^{-2}$ | 25 | 58 | $6 \times 10^{-8}$ | 190 | 4 | 2 |
| Cysteine | 121 | 100 | 3.78 | $1.53 \times 10^{-2}$ | 25 | 58 | $6 \times 10^{-8}$ | 190 | 4 | 2 |
| Leucine | 131 | 100 | 3.86 | $2.10 \times 10^{-2}$ | 25 | 58 | $6 \times 10^{-8}$ | 190 | 5 | 2 |
| Phenylalanine | 165 | 100 | 4.01 | $1.08 \times 10^{-2}$ | 25 | 58 | $6 \times 10^{-8}$ | 190 | 4 | 2 |
| Serine | 105 | 100 | 4.08 | $1.56 \times 10^{-2}$ | 58 | 111 | $6 \times 10^{-8}$ | 311 | 5 | 3 |
| Asparagine | 132 | 100 | 4.50 | $1.74 \times 10^{-2}$ | 58 | 111 | $1.4 \times 10^{-7}$ | 380 | 25 | 3 |
| Methionine | 149 | 100 | 3.90 | $1.35 \times 10^{-2}$ | 58 | 111 | $6 \times 10^{-8}$ | 311 | 5 | 3 |
| Tryptophan | 204 | 50 | 4.20 | $0.50 \times 10^{-2}$ | 58 | 111 | $6 \times 10^{-8}$ | 311 | 5 | 3 |
| Glutamine | 146 | 100 | 3.39 | $5.58 \times 10^{-2}$ | 57 | 115 | $3 \times 10^{-8}$ | 360 | 30 | 4 |

[a] Tip ID & OD are measured by SEM imaging.
[b] Chamber pressure is measured with an ion gauge.
[c] The tip number labels the four different nanopore ion sources used in the amino acid measurements.

#### 2 Water spectrum

Figure S1 shows the mass spectrum obtained from a control measurement of DI water sample with no amino acid present using a nanopore ion source with a 23 nm ID tip. The extraction voltage was 230 V and the emission current was 80 pA. A distribution of peaks corresponding to solvated hydronium ions ($H_3O^+(H_2O)_n$), with $n$ = 3 - 9 are visible in the spectrum. The most intense peak corresponds to $H_3O^+(H_2O)_5$ and the relative abundance decreases for larger clusters.

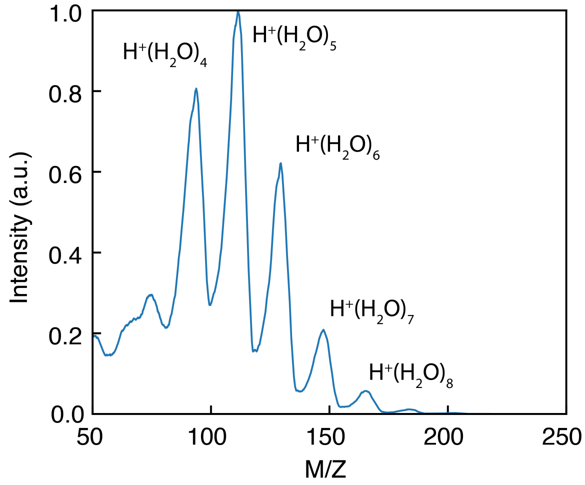


**Figure S1:** Mass spectrum of DI water obtained showing solvated hydronium ions.

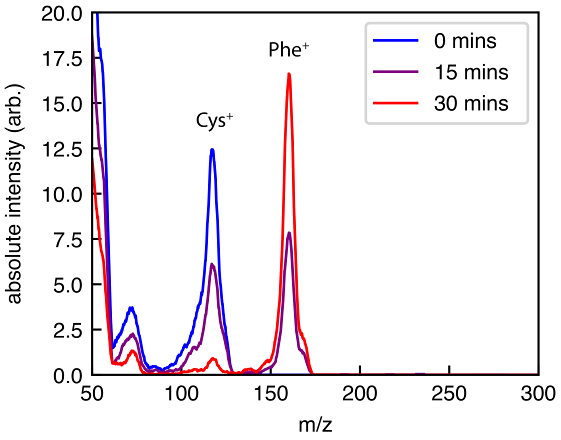


**Figure S2:** Mass spectra of obtained 0 (blue line), 15 (purple line), and 30 minutes (red line) following the introduction of 100 mM Phenylalanine into an ion source tip initially filled with 100 mM Cysteine.

### 3 Swapping solutions

Figure S2 shows three mass spectra obtained at 15 minute intervals while a 100 mM solution of cysteine was swapped for a 100 mM of phenylalanine. As the new solution gradually displaces the old solution at the tip of the ion source, peaks correponding to both amino acid types are visible in the spectrum.

### 4 Sensitivity and its relation to transmission efficiency

In this section we present an estimate of the minimum number of emitted analyte molecules required for a reliable detection in our MS experiments, which is an important measure of the sensitivity [1]. An alternative measure of sensitivity is the minimum detectable concentration of analyte, which we did not explore in this work. The quantity of ions emitted per quadrupole scan is given by $I_E$, integrated over the duration of a scan, $\Delta t$, and divided by the charge on each ion, $zq_e$,

$$N_{\text{emitted}} = \frac{I\Delta t}{zq_e}. \tag{S1}$$

Figure S3a-c shows mass spectra of a 100 mM aqueous arginine solution obtained with 1, 10, and 100 scans. The median scan duration is $\Delta t = 0.64$ s. A single scan (Fig. S3a) produces a spectrum composed of 55 individual ion detection events with a clear peak at $m/z = 175$. Thus, a single scan

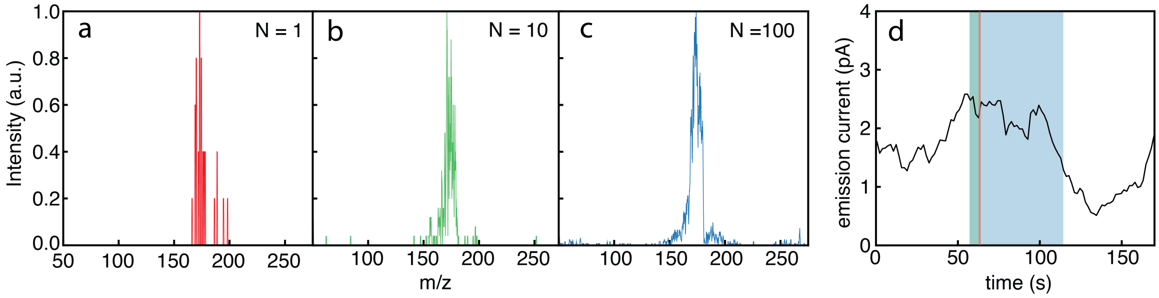


**Figure S3:** Mass spectra of 100 mM Arginine recorded with **(a)** 1 scan (≈ 0.6 seconds), **(b)** 10 scans (≈ 6 seconds), and **(c)** 100 scans (≈ 60 seconds). **d** Emission current as a function of time over the period in which the detections in **a, b** & **c** were measured. The red, green, and blue regions correspond to the 0.6 s, 6 s, and 60 s of data shown in **a, b** & **c**, respectively.

enables the identification of arginine. Fig. S3d plots $I_E$ during the measurement. Using Eq. (S1) with $I_E = 2.2$ pA gives $N_{\text{emitted}} = 8.8 \times 10^6$ analyte ions, or 15 attomoles. Note that $N_{\text{emitted}}$ provides an upper bound on the number of analyte molecules emitted because analyte only escapes in the form of ions, and non-analyte ions can carry some portion of the emission current. Using $N_{\text{emitted}}$ as we have done here thus gives a worst-case estimate of the amount of analyte consumed. The 15 attomole sensitivity of our measurement is similar to the typical sensitivity of a proteomics MS measurement, which is in the range of an attomole (∼1 million copies) to a femtomole (∼1 billion copies) [2]. However, its is worth noting that we are comparing measurements preformed on small amino acid ions with large proteins, which may differ in their emission characteristics.

Sensitivity depends on every step of the measurement leading up to detection, and any step could be the limiting one. ESI is often the limiting step, losing 99% or more of the analyte. In contrast, many ion detectors and mass analyzers (eg. orbitraps) waste very few ions. However, in our instrument, the quadrupole mass filter is the limiting step, because its narrow mass acceptance window rejects the vast majority of ions over the course of a scan [3]. For most of a scan, the acceptance window is far from the m/z of the analyte, and even when the two align, the transmission is typically about 10-20% or less [4, 5]. Additionally, a misalignment of the tip with respect to the central axis of the ion optics and the quadrupole of about 1 mm can cause significant deflection of the ion beam, reducing transmission through all subsequent parts of the instrument. We reason that significant improvements in sensitivity can be achieved by combining the nanopore ion source, which wastes very few ions, with a highly efficient ion trapping mass analyzer.

### 5 Simulations of ion and droplet trajectories through the magnetic sector

We preformed simulations to determine the $m/z$ range of particles expected to strike the Faraday cup and Faraday plate detectors used in the magnetic sector experiments. The trajectories of particles passing through the magnetic sector, which are subject to the Lorentz force, are calculated numerically by solving Newton's second law using a fourth-order Runge-Kutta scheme with a custom Python script. Particles are given a kinetic energy of $qV$, where $V = 500$ is the tip voltage used in the magnetic sector measurements presented in §2.4 of the main text, and $q$ is the charge on the particle. The magnetic field has a magnitude of 0.54 T, is oriented in the $+z$-direction within the bounds of the magnetic sector (a circle with a diameter of 6 cm), and vanishes outside these bounds. Given the geometry of the instrument, particles deflected less than 3.2° will strike the Faraday cup and contribute to $I_{\text{Drop}}$, while particles deflected by between 31° and 59° will strike the Faraday plate and contribute to ($I_{\text{Ion}}$). We find that particles with $75 < m/z < 315$ strike the Faraday plate, while particles with $m/z > 33,000$ strike the Faraday cup. The latter $m/z$ value corresponds to a 14 nm radius water droplet charged to the Rayleigh limit [6] as well as a 7 nm radius droplet sufficiently charged to readily evaporate ions [7].

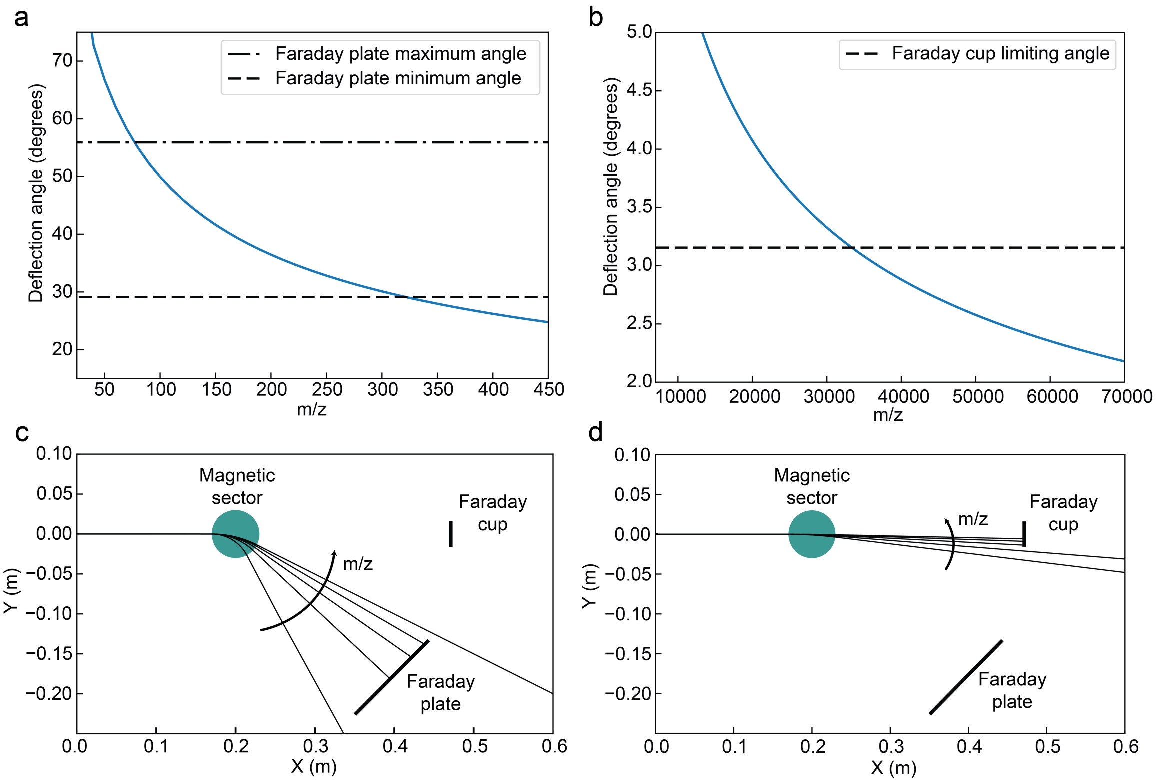


**Figure S4:** **a** Simulation results of deflection angle vs $m/z$ for small ions. The region between the dashed lines represents the position and extent of the Faraday plate detector, so ions with $70 < m/z < 325$ should strike the Faraday plate. **b** Simulation results of deflection angle vs $m/z$ for nanoscale charged droplets. The region below the dashed line represents the position and extent of the Faraday cup aperture, so species with $m/z > 33000$ be detected. **c** Selection of simulated trajectories for five singly charged ions of mass between 60 and 460 amu, the arrow points in the direction of increasing $m/z$, and the blue circle represents the magnetic sector. **d** Selection of simulated trajectories for five nanodroplets charged to the Rayliegh limit with radius between 5 and 25 nm, the arrow points in direction of increasing $m/z$.

### 6 Non-normalized mass spectra from Fig. 4

Figure S5 shows the same mass spectra shown in Fig. 4 of the main text without any normalization. While the pH-dependent change in the ratio of amino acid signal to $Na^+$ signal is still apparent, there is also a global increase in signal at higher pH in both the histidine and arginine data. This increase in signal at the higher pH value was mirrored by an increase in the measured emission current from about 2 pA to 5 pA for the histidine measurement, and from about 2 pA to 20 pA for the arginine measurement. We believe this pH-dependence in the overall emission current may be due to changes in the surface charge density of the glass capillary which governs surface conduction [8, 9].

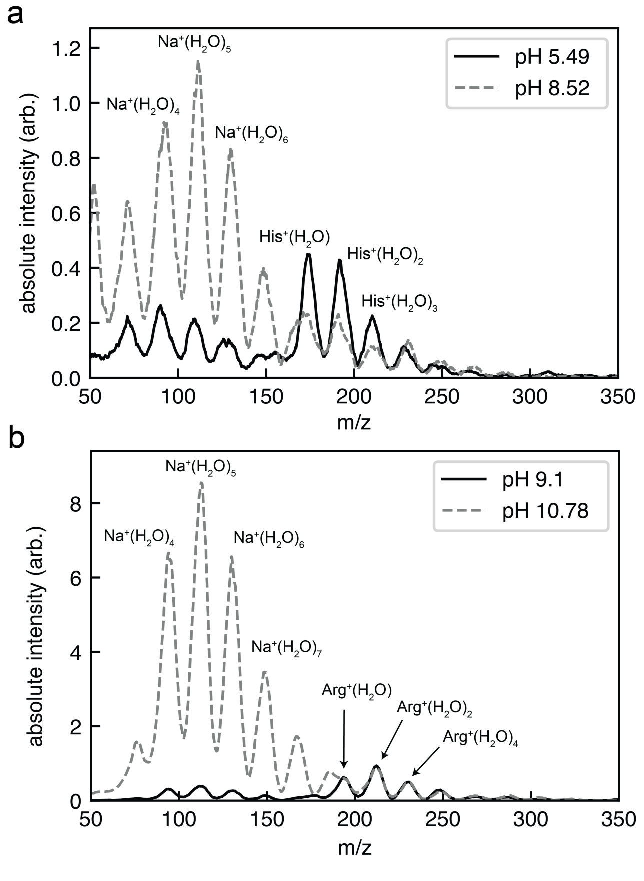


**Figure S5:** Unnormalized mass spectra from Fig. 4 in the main text.

#### 7 Probability that an emitted ion collides with a gas molecule

In conventional ESI, collisions with gas molecules are the primary mechanism that removes solvent molecules from ions. However, our instrument operates under high vacuum conditions where collisions with gas molecules are rare. We applied the kinetic theory of gases to calculate the probability that an emitted ion cluster would collide with at least one gas molecule. This addresses the question of whether ions emerge from the solution unsolvated, or whether they emerge with a solvation shell which is knocked off by collisions with gas molecules on their way to the detector.

We take the distribution of gas particles within the vacuum chamber to be the sum of two components, a uniform background of number density $n_b$ and a distribution of water molecules $n_w$ that evaporate from the meniscus at the nanopore tip. The background gas pressure in our experiments was typically around $7 \times 10^{-8}$ torr (see table S1). The mean free path of a water molecule in nitrogen gas at this pressure is greater than 1 km, so we model the evaporating water molecules as travelling radially outward from a hemispherical meniscus, as illustrated in Fig. S1a. The density of water molecules decays as $n_w \sim r^{-2}$ near the ion source, where $r$ is the distance from the center of the hemisphere. The rate at which water evaporates from the nanoscale liquid meniscus into vacuum is difficult to establish precisely [10]. However, the flux of water molecules evaporating from the liquid surface at a given temperature cannot exceed the flux of incoming molecules at equilibrium. Thus, if we subtract the inward-moving molecules, the highest possible density that the (outward-moving) evaporating water molecules could achieve on the vacuum side of the meniscus is half the water vapor density at equilibrium. We therefore estimate the upper limit of the probability that an emitted ion collides with a gas molecule by assuming the density of water vapor on the vacuum side of the meniscus is $\frac{p_0}{2k_BT}$, where $p_0 = 17.5$ torr is the approximate equilibrium vapor pressure of water at 20 °C [11], and $k_BT$ is the thermal energy.

We are interested in the cumulative probability that an ion cluster leaving the source undergoes at least one collision before reaching $r$, $C_P(r)$. To compute $C_P(r)$, it is easier to first consider the probability that an ion survives to a distance $r$ without undergoing a collision, $P_s(r) = 1 - C_P(r)$, which can be defined recursively as

$$P_s(r) = P_s(r - dr)\left(1 - P_c(r - dr \to r)\right), \tag{S2}$$

where $P_c\left(r - dr \to r\right)$ is the probability that a collision occurs between $r - dr$ and $r$. In the limit of dilute gasses and infinitesimal displacements,

$$P_c(r - dr \to r) = \sigma_w n_{w,0} \left(\frac{r_0}{r}\right)^2 dr + \sigma_b n_b dr, \tag{S3}$$

where $\sigma_w$ is the cross section for a collision between an ion cluster and a water molecule, $\sigma_b$ is the cross section for an ion collision with a background gas molecule, $n_{w,0}$ is the number density of water on the vacuum side of the meniscus, and $r_0$ is the radius of the meniscus. Combining Eqs. (S2) and (S3) gives a differential equation for $P_s$

$$\frac{P_s(r) - P_s(r - dr)}{P_s(r - dr)} = \frac{dP_s(r)}{P_s(r)} = -\sigma_w n_{w,0} \left(\frac{r_0}{r}\right)^2 dr - \sigma_b n_b dr. \tag{S4}$$

Integrating Eq. (S4) from $r_0$ to $r$, and applying the boundary condition $P_s(r_0) = 1$ gives

$$P_s(r) = \exp\left[-\sigma_w n_0 \left(\frac{1}{r_0} - \frac{1}{r}\right) - \sigma_b n_b (r - r_0)\right]. \tag{S5}$$

Finally, the cumulative probability that an ion experiences at least one collision before reaching $r$ is given by $1 - P_s(r)$,

$$C_P(r) = 1 - \exp\left[-\sigma_w n_0 \left(\frac{1}{r_0} - \frac{1}{r}\right) - \sigma_b n_b (r - r_0)\right]. \tag{S6}$$

Figure S1b plots $C_P(r)$ using parameters corresponding to the conditions of our MS measurements. The cross sections $\sigma_w$ and $\sigma_b$ are given respectively by $\pi(a_i + a_w)^2$ and $\pi(a_i + a_b)^2$, where $a_w = 1.3$ Å is the kinetic radius of water, $a_b = 1.8$ Å is the kinetic radius of the background gas molecules [12], and $a_i = 7$ Å is the approximate radius of an amino acid with a full hydration shell, which we estimated using the Stokes-Einstein relation and reported diffusion coefficients of amino acids in water [13]. $n_b = 2.25 \times 10^{15}$ m$^{-3}$ and $n_{w,0} = 6.44 \times 10^{23}$ are the number densities of background gas and water molecules on the vacuum side of the meniscus, respectively, at a background pressure of $7 \times 10^{-8}$ torr and equilibrium water vapor pressure of 17.5 torr. We use a meniscus radius of $r_0 = 30$ nm. Figure S1b shows $C_P(r)$ rise rapidly with $r$ over the first 100 nm before saturating at about 1.8 %. $C_P(r)$ increases slowly over centimeter-scale distances due to the finite density of background gas molecules. In total, $C_P(r)$ rises to 2.1% over the 50 cm distance from the ion source to the detector.

These results show that, in contrast to conventional electrospray, collisions between gas particles and ion clusters emitted from the nanopore ion source are rare. The majority of ions follow collisionless trajectories from the source to the detector because the instrument operates under high vacuum conditions. The lack of collisions implies that ions are emitted in the same state as when they are detected, leading us to conclude that the nanopore ion source is capable of emitting predominantly desolvated amino acid and small peptide ions.

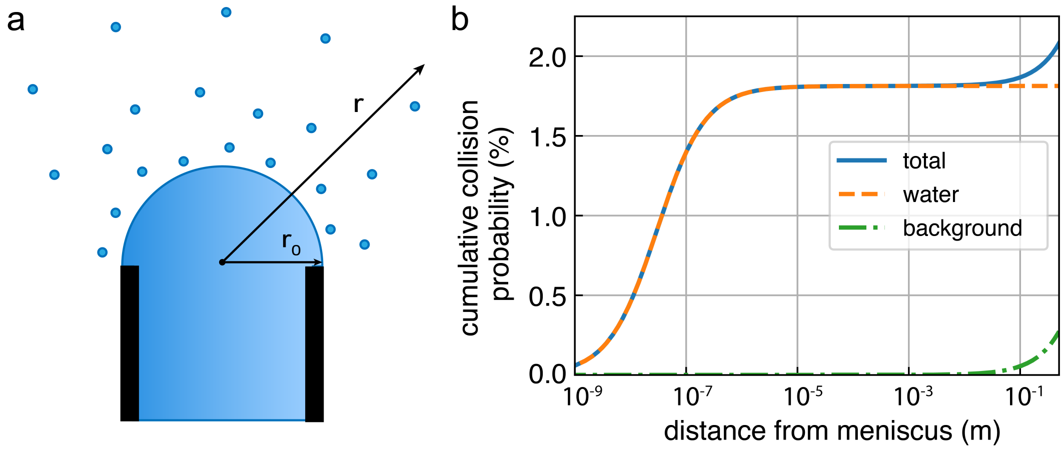


**Figure S6: a.** Illustration of water evaporating from a hemispherical meniscus showing the tip radius $r_0$ and the distance $r$. **b.** Plot of $C_P(r)$ (solid blue line) and the separate contributions from water molecules (dashed orange line) and background gas (dash-dot green line) to the total probability of undergoing at least one collision. The orange dashed curve shows the contribution from evaporated water molecules, the green dashed and dotted curve shows the contribution from the background gas, and the blue solid curve shows the total collision probability obtained by summing the contributions from the water and background gas. The calculation used $n_b = 2.25 \times 10^{15}$ m$^{-3}$, $n_{w0} = 6.44 \times 10^{23}$ m$^{-3}$, $a_w = 1.3$ Å, $a_b = 1.8$ Å, $a_i = 7$ Å, and $r_0 = 30$ nm. The plots extend to $r = 0.5$ m, the total distance from the meniscus to the detector.
